# SupportNet: a novel incremental learning framework through deep learning and support data

**DOI:** 10.1101/317578

**Authors:** Yu Li, Zhongxiao Li, Lizhong Ding, Yuhui Hu, Wei Chen, Xin Gao

## Abstract

**Motivation:** In most biological data sets, the amount of data is regularly growing and the number of classes is continuously increasing. To deal with the new data from the new classes, one approach is to train a classification model, e.g., a deep learning model, from scratch based on both old and new data. This approach is highly computationally costly and the extracted features are likely very different from the ones extracted by the model trained on the old data alone, which leads to poor model robustness. Another approach is to fine tune the trained model from the old data on the new data. However, this approach often does not have the ability to learn new knowledge without forgetting the previously learned knowledge, which is known as the *catastrophic forgetting* problem. To our knowledge, this problem has not been studied in the field of bioinformatics despite its existence in many bioinformatic problems.

**Results:** Here we propose a novel method, SupportNet, to solve the catastrophic forgetting problem efficiently and effectively. SupportNet combines the strength of deep learning and support vector machine (SVM), where SVM is used to identify the support data from the old data, which are fed to the deep learning model together with the new data for further training so that the model can review the essential information of the old data when learning the new information. Two powerful consolidation regularizers are applied to ensure the robustness of the learned model. Comprehensive experiments on various tasks, including enzyme function prediction, subcellular structure classification and breast tumor classification, show that SupportNet drastically outperforms the state-of-the-art incremental learning methods and reaches similar performance as the deep learning model trained from scratch on both old and new data.

**Availability:** Our program is accessible at: https://github.com/lykaust15/SupportNet.

## 1 INTRODUCTION

Since the breakthrough in 2012 (Krizhevsky *et al.*, 2012), deep learning has achieved great success in various fields (LeCun *et al.*, 2015; Silver *et al.*, 2016; Sutskever *et al.*, 2014; He *et al.*, 2016). It has also facilitated the development of bioinformatics greatly (Min *et al.*, 2017; Alipanahi *et al.*, 2015; Li *et al.*, 2018b; Dai *et al.*, 2017). However, despite its impressive achievements, in addition to the weak theoretical support (Brutzkus *et al.*, 2017; Brutzkus and Globerson, 2017), there are still several bottlenecks related to the practical part of deep learning waiting to be solved, such as adversarial attack (Papernot *et al.*, 2016), lacking interpretability (Lipton, 2016), catastrophic forgetting (Kemker *et al.*, 2017), and failure to model uncertainty (Gal and Ghahramani, 2016). Among them, *catastrophic forgetting* means that a well-trained deep learning model tends to completely forget all the previously learned information when learning new information (McCloskey and Cohen, 1989). That is, once a deep learning model is trained to perform a specific task, it cannot be trained easily to perform a new similar task without affecting the original task’s performance dramatically. For example, suppose after we have trained a deep learning model which can recognize 1000 flower species based on the given flower pictures, the data of the 1001st flower species appear. If we only train the model with the new coming data, the model’s performance on classifying the previous 1000 species would be unacceptable, even worse than random guess (Rebuffi *et al.*, 2016). In other words, deep learning models, unlike human and animals, do not have the ability to continuously learn over time and different datasets by incorporating the new information while retaining the previously learned experience, which is known as *incremental learning*.

The problem is aggravated in the bioinformatics field due to the explosion of biological data. In the past decade, we have witnessed the dramatic increase in the amount of the genomic data (Marx, 2013), protein sequence data (UniProt, 2007) and the emergence of various databases (Zou *et al.*, 2015). As a natural consequence of data accumulation, the number of classes within each dataset is also increasing. For example, the label space of the EC system (Webb *et al.*, 1992) is continuously expanding; the number of entries of ontology (Ashburner *et al.*, 2000) is also constantly increasing; new plant species (Joppa *et al.*, 2011) are being found along the time as well. With the new coming data of new classes, the previously well-trained deep learning model for enzyme function prediction (Li *et al.*, 2018a), plant species recognition (Lee *et al.*, 2015), and disease prediction (Mohanty *et al.*, 2016; Chen *et al.*, 2017) would face the serious problem of catastrophic forgetting. In spite of the severity of the problem, currently, to our knowledge, this problem has not been studied in the bioinformatics field.

On the other hand, human and other animals have shown significant superiority over artificial intelligence systems in dealing with catastrophic forgetting and incorporating new knowledge with little, if any, negative effect on the previously learned knowledge (Bremner *et al.*, 2012). Two major theories have been proposed to explain this ability to perform incremental learning. The first theory is Hebbian learning (Hebb, 1949) with homeostatic plasticity (Zenke *et al.*, 2017), which focuses on the mechanism of neurosynaptic plasticity regulating the stability-plasticity balance in the brain. During the early stage of human development, the human brain has a very high degree of plasticity, which enables the brain to learn knowledge by changing the synaptic strength and building new connections (Hensch *et al.*, 1998). After those critical periods, although a certain degree of plasticity would be preserved for the brain reorganization, the rate of synaptic plasticity would decrease so that the previously learned information would be protected against the interference when learning new tasks (Cichon and Gan, 2015). The second theory is the complementary learning system (CLS) theory (Mcclelland *et al.*, 1995; OReilly *et al.*, 2014), which explains how human beings extract high-level structural information while retaining episodic memories. Specifically, the CLS theory suggests that hippocampus stores episodic memory, enabling fast learning of arbitrary information while the neocortex would store the structured knowledge. The two different brain areas are connected for memory storage and retrieval. This theory suggests that by separating the two different memories into different areas, the brain can protect the consolidated knowledge, although the detailed mechanism is waiting to be elucidated.

As for neural network systems, the most straightforward and pragmatic method to avoid catastrophic forgetting is to retrain a deep learning model completely from scratch with all the old data and new data (Parisi *et al.*, 2018). However, this method is proved to be very inefficient (Parisi *et al.*, 2018). Moreover, the new model learned from scratch may share very low similarity with the old one, which results in poor learning robustness. Inspired by the above two major neurophysiological theories of human incremental learning, researchers have proposed three main categories of neural network systems to alleviate the effect of catastrophic forgetting. The first category is the regularization approach (Kirkpatrick *et al.*, 2017; Li and Hoiem, 2016; Jung *et al.*, 2016), which is inspired by the plasticity theory (Benna and Fusi, 2016). The core idea of such methods is to incorporate the plasticity information of the neural network model into the loss function so that to prevent the parameters from varying significantly when learning new information. These approaches are proved to be able to protect the consolidated knowledge (Kemker *et al.*, 2017). However, due to the fixed size of the neural network, there is a trade-off between the performance of the old and new tasks (Kemker *et al.*, 2017). The second class uses dynamic neural network architectures (Rebuffi *et al.*, 2016; Rusu *et al.*, 2016; Lopez-Paz and Ranzato, 2017). To accommodate the new knowledge, these methods dynamically allocate neural resources or retrain the model with an increasing number of neurons or layers. Intuitively, these approaches can prevent catastrophic forgetting but may also lead to scalability and generalization issues due to the increasing complexity of the network (Parisi *et al.*, 2018). The last category utilizes the dual-memory learning system, which is inspired by the CLS theory (Hinton and Plaut, 1987; Lopez-Paz and Ranzato, 2017; Gepperth and Karaoguz, 2016). Most of these systems either use dual weights or take advantage of pseudo-rehearsal, which draw training samples from a generative model and replay them to the model when training with new data. However, how to build an effective generative model remains a difficult problem.

Despite the development on tackling this problem, existing approaches cannot be applied to bioinformatics data directly due to the differences in the nature and properties of the data, which will be shown in Section 4. Here, we propose a novel method, inspired by the above two neurophysiology theories and the intrinsic sparsity of support vector learning, to perform class incremental deep learning efficiently when encountering data from new classes for bioinformatic tasks (Fig. 1). Our method maintains a support dataset for each old class, which is much smaller than the original dataset of that class, and shows the support datasets to the deep learning model every time there is a new class coming in so that the model can “review” the representatives of the old classes while learning new information. The idea of showing representative old data to the new model is known as *rehearsal*, which was proposed recently in the computer vision field (Rebuffi *et al.*, 2016). However, our method is innovative in the sense that we select the support dataset through a novel combination of deep learning and support vector machine (SVM). Using deep learning to extract high level features and SVM to detect the support vectors, we are more likely to construct the support data which are of vital importance for the classification. Besides, following the idea of the Hebbian learning theory, to reduce the plasticity of the deep learning model, we utilize two *consolidation regularizers*, which constrain the deep learning model to produce similar representation for old and new data, and retain the performance at the same time. Furthermore, to extract better high level representations of the original data, which are crucial for rehearsal-based methods, we incorporate squeeze-and-excitation network (SENet) (Hu *et al.*, 2017), considering not only the spatial information but also the channel information, as the feature extractor component in our method, which is usually omitted by other convolutional models. In summary, this paper has the following main contributions:

- We propose the first method to perform the class incremental learning in the bioinformatics field, which alleviates the notorious catastrophic forgetting problem when using deep learning to investigate biological data.
- We propose a novel way of selecting support data through the combination of deep learning and SVM.
- We propose a novel regularizer, namely, *feature regularizer*, which stabilizes the deep learning network and maintains the high level feature representation of the old information.

**Fig. 1.**
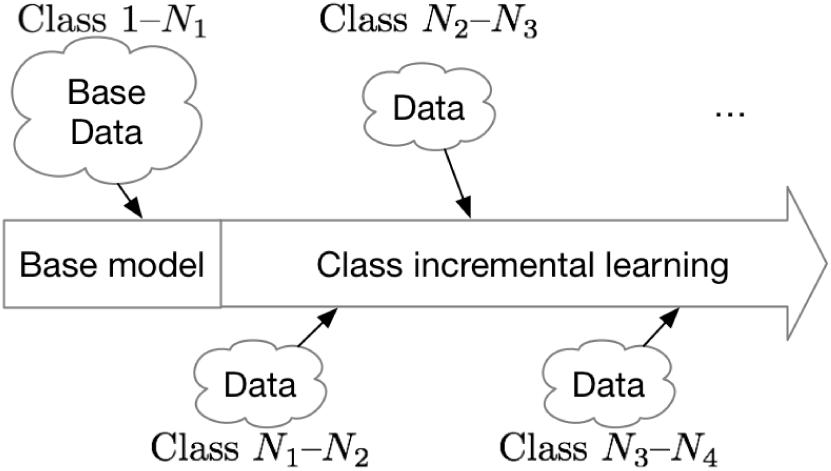
Illustration of class incremental learning. After we train a base model using all the available data at a certain time point (e.g., classes 1, …, *N*_1_), the new data belonging to new classes may continuously appear (e.g., classes *N*_1_, …, *N*_2_, classes *N*_2_, …, *N*_3_, etc.), which is a commonly seen scenario in biology.

## 2 RELATED WORKS

### 2.1 EWC

Elastic weight consolidation (EWC) (Kirkpatrick *et al.*, 2017), inspired by the synaptic plasticity theory, is a very practical solution to solve the catastrophic forgetting problem when training a sequential set of classification models. By considering the Fisher information of each weight and adding a penalty term to the loss function, this method prevents weights from changing too much if the weights are closely related to the classifiers on the old data. Slowing down the learning of the task-related weights, EWC can retain the learned knowledge when incorporating new information, which makes it suitable for incremental supervised learning and reinforcement learning (Parisi *et al.*, 2018). Despite its additional computational cost and limited applications to the low-dimensional output space, EWC was shown to be a well-recognized method for solving the catastrophic forgetting problem in deep learning (Parisi *et al.*, 2018; Kemker *et al.*, 2017).

### 2.2 iCaRL

iCaRL (Rebuffi *et al.*, 2016) is currently the state-of-the-art method for class incremental learning in the computer vision field. It combines deep learning with *k* nearest neighbor (KNN), using deep learning to extract the high level feature representation for each data point and deploying KNN as the final classifier. During classification, it computes the average data representation of a certain class using all the training data (or preserved examplars) belonging to that class, finds the nearest class-averaged representation for the test data, and assigns the class label accordingly. In order to reduce the memory footprint when the number of class increases dramatically, the approach maintains an examplar set for each class. To construct the examplars, it chooses those data points which are closest to the averaged representation of that class. By combining old and new data, it avoids catastrophic forgetting and achieves the best performance on the commonly used benchmark datasets in the computer vision field (Rebuffi *et al.*, 2016; Parisi *et al.*, 2018). Despite its impressive performance on those datasets, the power of the method degrades drastically on bioinformatics datasets, which will be shown in Section 4.

## 3 METHODS

Due to the highly complex and stochastic environment, biological systems are usually of high variability, which makes most kinds of biological data noisy in nature. Besides, several kinds of biological data, such as biomedical images and gene expression data, can be of high dimensionality. Furthermore, sometimes the data, such as the enzyme function data and the gene ontology data, may also have enormous label space which consists of a large number of classes. Taking those properties into consideration, we design the following framework to perform the class incremental learning for biological data (Fig. 2). In addition to the deep learning model, which is used to extract the high level features from the original noisy and high-dimensional inputs (Section 3.3), the two novel components of our framework are support data selector (Section 3.1) and consolidation regularizers (Section 3.2). Training an SVM using the high level features extracted by the SENet feature extractor, we can detect which data points are important for the classification based on the support vector information and thus select the support data for each class, which will be shown to the deep learning model for future training to prevent the network from catastrophic forgetting. Compared to training a new model from scratch, this approach is much less computationally costly and requires much less memory. On the other hand, because of the further training, the high level features may change, which invalidates the support data, and thus decreases the performance of the deep learning model on the old classes. To prevent such deterioration, we add two consolidation regularizers into the loss function. The first one is the *feature regularizer*, which is applied to the high level feature layer, ensuring the extracted features of the old classes remain consistent and thus guaranteeing the effectiveness of the support data for the old classes. The second one is the *EWC regularizer*, which consolidates the weights critical for older classes and moves learned parameters to the region where both the old data and the new data have low loss.

**Fig. 2.**
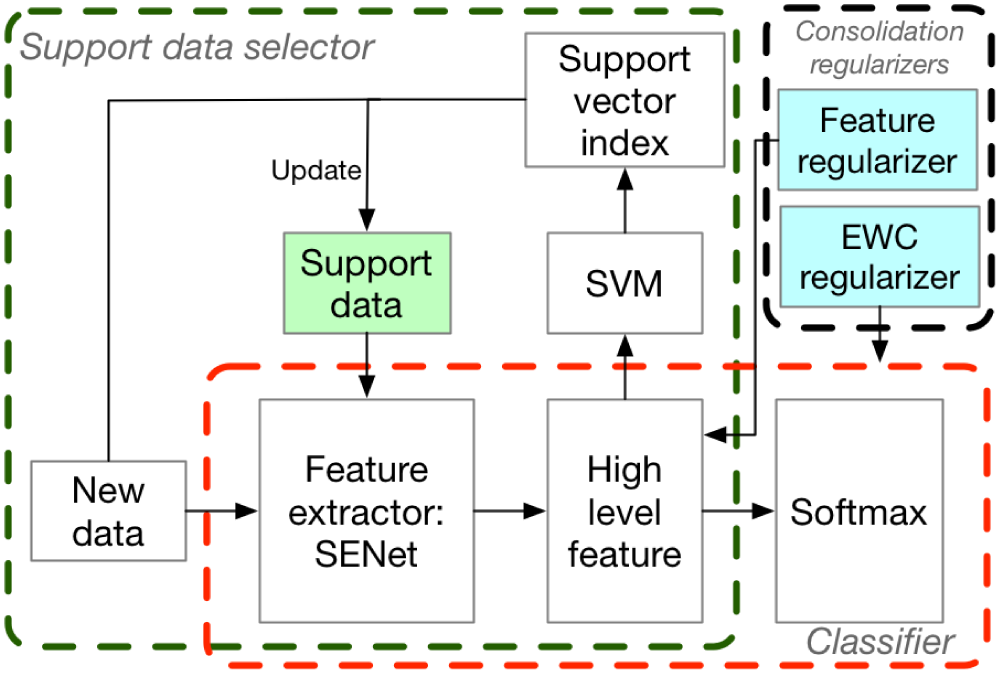
Overview of our framework. The core idea is to incrementally train the deep learning model efficiently using the new class data and the support data of the old classes, regularized by consolidation regularizers, so that the trained model is free of catastrophic forgetting and can classify all the observed classes well. The two vital components are the support data selector (the green dashed box) and the two consolidation regularizers (the black dashed box). The selector takes advantage of the feature extraction power of deep learning and the intrinsic sparsity of SVM, and is able to choose a small while representative support dataset. The two consolidation regularizers consolidate the old information in the network by forcing the deep learning model to produce consistent representation for old data (the feature regularizer) and to reduce the plasticity of the critical weights for old classes (the EWC regularizer).

### 3.1 Support Data Selector

According to (Sirois *et al.*, 2008; Pallier *et al.*, 2003), even human beings, who are proficient in incremental learning, could not deal with catastrophic forgetting perfectly. On the other hand, a common strategy for human beings to overcome forgetting during learning is to review the old knowledge frequently (Murre and Dros, 2015). Actually, during reviewing, we usually do not review all the details, but rather the important ones, which are often enough for us to grasp the knowledge. Inspired by this, we design the support dataset and the review training process. During incremental learning, we maintain a support dataset for each class, which is fed to the model together with the new data of the new classes. In other words, we want the model to review the representatives of the previous classes when learning new information.

The main question is thus how to build an effective support data selector to construct such support data. Inspired by the sparsity of support vector learning, we design a novel support data selection process which is based on the support vectors of SVM. After obtaining the high level feature representations of the original input using SENet, we train an SVM classifier with these features, which can be considered as an approximation of the last layer of the deep learning model. By performing the SVM training, we detect the support vectors, which are of crucial importance for the classification. We define the original data which correspond to these support vectors as the *support data candidates*. If the required number of support data is smaller than that of the support vectors, we will sample support data candidates to obtain the required number. Fig. 3 summarizes the idea of selecting the support data.

**Fig. 3.**
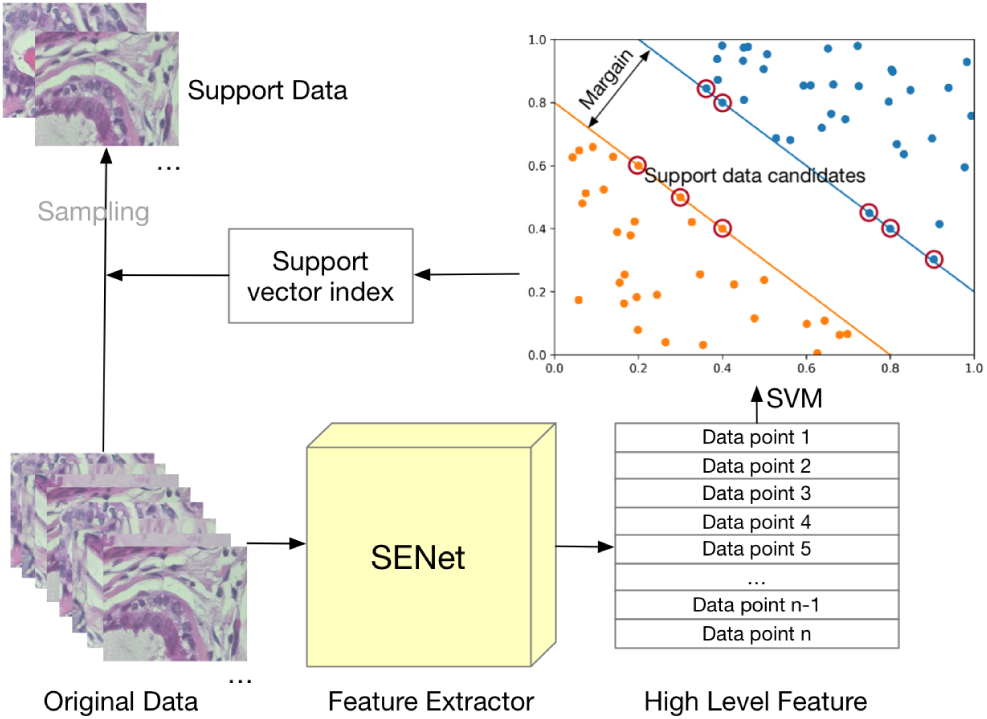
Support data selector. We first feed the data in the original feature space to the SENet module, which extracts high level features. We then use an SVM on these features to approximate the classification layer of the deep learning model. After detecting the support vectors from SVM, we can find the original data corresponding to these support vectors, which are then used to construct and update the support data.

Note that SVM here is only used to select the support data candidates, but not to be the final classifier. This is due to the fact that SVM is not as powerful as deep learning on multi-class classification, especially when the number of classes is large.

### 3.2 Consolidation Regularizers

Although the support data and the review training process largely alleviate the catastrophic forgetting problem, the model may still be subject to it if we only use that technique, due to two reasons. Firstly, since the support data selection depends on the high level features produced by SENet, which are fine tuned on new data, the old data feature representations may change over time. As a result, the previous support vectors for the old data may no longer be support vectors for the new data, which makes the support data invalid. Secondly, because the deep learning model has very high expressibility (Brutzkus *et al.*, 2017) while the support data are often of limited size, the model is very likely to become overfitting. To solve these issues, we add two consolidation regularizers to consolidate the learned knowledge: the feature regularizer, which forces the model to produce fixed representation for the old data over time, and the EWC regularizer, which consolidates the weights contributing to the old class classification significantly into the loss function.

#### 3.2.1 Feature Regularizer

According to Fig. 3, the selection of the support data largely depends on the support vectors of the SVM classifier, which is further determined by the SENet since the output of the SENet is used to train the SVM. If the high level representations of the old support data, produced by the SENet, change over time, the previous support vectors may no longer be the support vectors if we retrain an SVM classifier, which makes the constructed support data no longer valid. To avoid this issue, we add the following feature regularizer into the loss function to force the SENet to produce fixed representation for old data.

Suppose *g*(*x*; *θ*) is the feature representation produced by the SENet parameterized by *θ* for the input *x*, the feature regularizer is defined as follows:

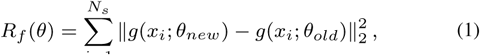

where *θ new* is the parameters for the SENet trained with the support data from the old classes and the new data from the new class(es); *θ_old_* is the parameters for the SENet of the old data; and *N_s_* is the number of support data.

This regularizer requires us to preserve the feature representation produced by the SENet for each support data, which could lead to potential memory overhead. However, since it operates on a very high level representation, which is of much less dimensionality than the original input, the overhead is neglectable.

#### 3.2.2 EWC Regularizer

According to the Hebbian learning theory, after learning, the related synaptic strength and connectivity are enhanced while the degree of plasticity decreases to protect the learned knowledge. Guided by this neurophysiological theory, the EWC regularizer (Kirkpatrick *et al.*, 2017) was designed to consolidate the old information while learning new knowledge. The core idea of this regularizer is to constrain those parameters which contribute significantly to the classification of the old data. Specifically, the more a certain parameter contributes to the previous classification, the harder constrain we apply to it to make it unlikely to be changed. That is, we make those parameters that are closely related to the previous classification less “plastic”. In order to achieve this goal, we calculate the Fisher information for each parameter, which measures its contribution to the final prediction, and apply the regularier accordingly.

Formally, the Fisher information for the parameters *θ* can be calculated as follows:

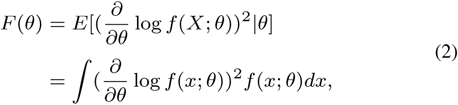

where *f* (*x*; *θ*) is the functional mapping of the entire deep learning model.

The EWC regularizer is defined as follows:

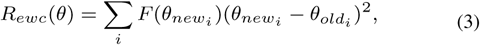

where *i* iterates all the parameters of the model.

There are two major benefits of using the EWC regularizer in our framework. Firstly, the EWC regularizer reduces the “plasticity” of the parameters that are important to the old classes and thus guarantees stable performance over the old classes. Secondly, by reducing the capacity of the deep learning model, the EWC regularizer prevents overfitting to a certain degree. The function of the EWC regularizer could be considered as changing the learning trajectory pointing to the region where the loss is low for both the old and new data, which is illustrated in Fig. 4.

**Fig. 4.**
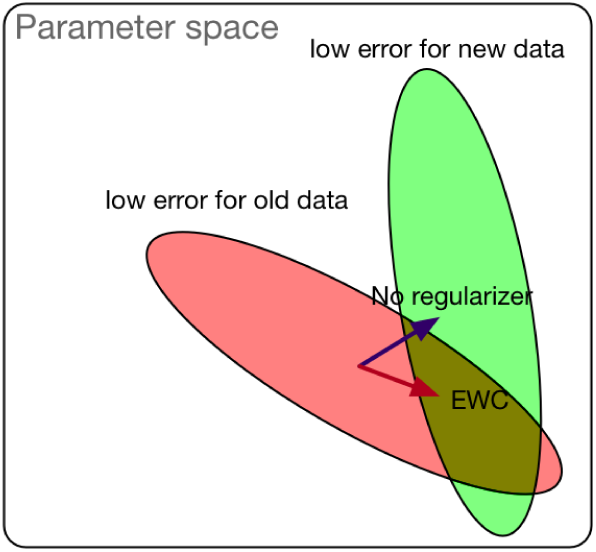
Illustration of the EWC regularizer. In the parameter space, the parameter sets which have low error for the old data (orange oval) and for the new data (gray oval) are not the same, but often overlap because the old and new data are related. If we do not add any regularizer, or only add the L1 or L2 regularizer, which does not have the capability of retaining old information, the learned parameters are likely to move to the region that is good for the new data, and thus the error is high for the old data. In contrast, the EWC regularizer pushes the learning to the overlapping region.

#### 3.2.3 Loss Function

The loss function of a traditional deep learning model is the cross entropy loss, which is defined as follows:

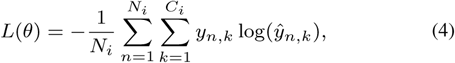

where *N_i_* is the training data size, including the new data and the support data, for the *i*th training round; *C_i_* is the total number of classes for the *i*th training round; *y_n_*,_*k*_ is the ground truth of the nth data sample belonging to class *k*; *ŷ_n_*,_*k*_ is the corresponding predicted probability.

After adding the feature regularizer and the EWC regularier, the loss function becomes:

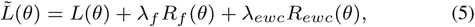

where *λ_f_* and *λ_ewc_* are the coefficients for the feature regularizer and the EWC regularizer, respectively.

After plugging Eq. (1), (3) and (4) into Eq. (5), we obtain the regularized loss function:

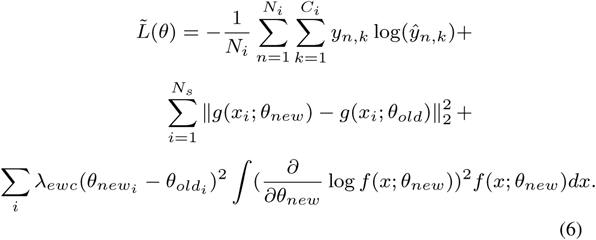

### 3.3 Feature Extractor

Similar to other deep learning methods, in which the feature representation learning is of vital importance, the feature extractor in our framework can also affect the performance significantly. Among numerous studies of exploring the deep learning model architectures, many of them focus on investigating the input’s spatial information with various filters and connections, such as the AlexNet (Krizhevsky *et al.*, 2012), VGG (Simonyan and Zisserman, 2014), ResNet (He *et al.*, 2016), ResNext (Xie *et al.*, 2016), and GoogLeNet (Szegedy *et al.*, 2014). In contrast, SENet (Hu *et al.*, 2017), in addition to utilizing the spatial information with 2D filters, further explores the information hidden in different channels by learning weighted feature maps from the initial convolutional output. The main idea of the residual network, on which the SENet is based, is to utilize a traditional convolutional layer within the residual block, which consists of the convolutional layer and a shortcut of the input, to model the residual between the output feature maps and the input feature maps. Despite the impressive performance of the residual block, it cannot explore the relation between different channels of the convolutional layer output. To overcome this issue, the SENet consummates the residual block with additional components which learn scale factors for different channels of the intermediate output and rescale the values of those channels accordingly. Intuitively, the traditional residual network considers different channels equally while the SENet takes the weighted channels into consideration. Using SENet as the feature extractor, which considers both the spatial information and the channel information, we are more likely to obtain a good structured high level representation of the original input data, which is vital for the support data selection process and the downstream classification task. Fig. 5 illustrates the main difference between the residual block and the SENet block.

**Fig. 5.**
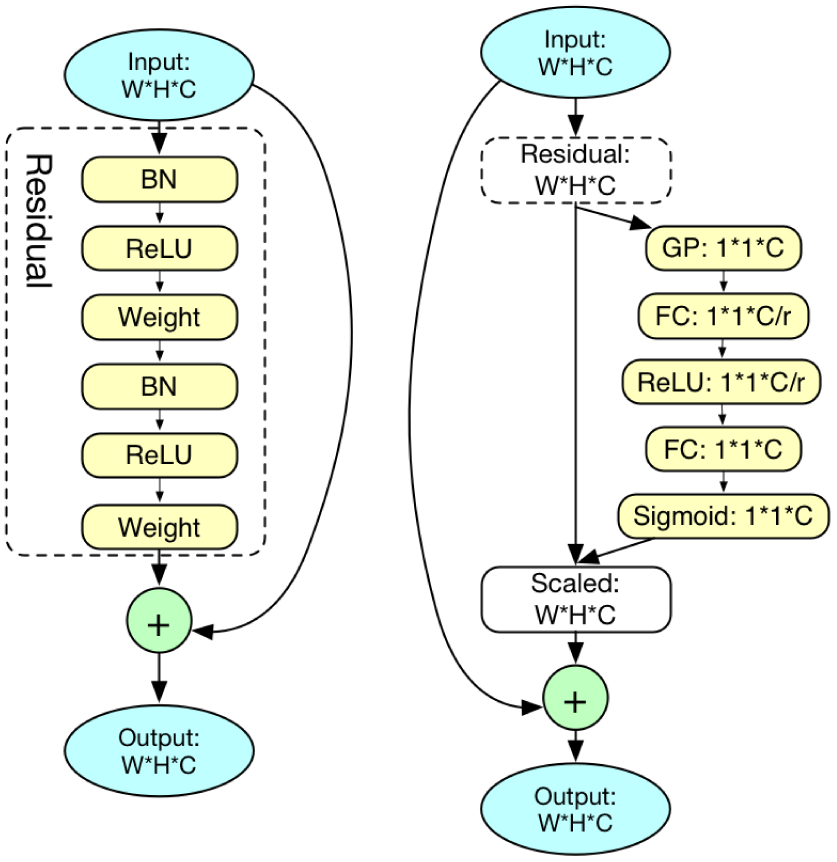
Comparison between the residual block (left) and the SENet block (right). In the residual block, the input feature maps, with dimensionality as *W* (weight) by *H* (height) by *C* (channels), go through two ‘BN’ (batch normalization) layers, two ‘ReLU’ activation layers and two ‘weight’ (linear convolution) layers. The output of these six layers is added to the original input feature maps element-wisely to obtain the residual block output feature maps. The SENet block extends the residual block by considering the channel information. After obtaining the residual layer output, it does not add the output directly to the original input. Instead, it learns a scaling factor for each channel and scales the channels accordingly, after which the scaled feature maps are added to the input element-wisely to obtain the SENet block output. To learn the scale vector, it first applies a ‘GP’ (global average pooling) layer onto the residual layer output, whose dimensionality is *W* by *H* by *C*, to obtain a vector with length *C*. After that, two ‘FC’ (fully connected) layers with ReLU and Sigmoid activation functions are used respectively to learn the final scaling vector. The hyper-parameter ‘r’, which determines the number of nodes in the first fully connected layer, is usually set as 16. By considering both the spatial information and the channel information comprehensively, the SENet is more likely to learn a better high level representation of the original input (Hu *et al.*, 2017).

### 3.4 SupportNet

Combining the deep learning model, which consists of the SENet feature extractor and the final fully connected classification layer, the novel support data selector, and the two consolidation regularizers together, we propose a highly effective framework, SupportNet (Fig. 2), which can perform class incremental learning for biological data without catastrophic forgetting. Our framework can resolve the catastrophic forgetting issue in two ways. Firstly, the support data can help the model to review the old information during future training. Despite the small size of the support data, they can preserve the distribution of the old data quite well, which will be shown in Section 4.7. Secondly, the two consolidation regularizers consolidate the high level representation of the old data and reduce the plasticity of those weights, which are of vital importance for the old classes.

## 4 RESULTS

### 4.1 Datasets

#### 4.1.1 Enzyme Function Prediction Dataset

This dataset^1^ is from our previous work (Li *et al.*, 2018a), which predicts the function of enzymes from their sequences through a novel deep learning architecture. After preprocessing the entire SWISS-PROT (Bairoch and Apweiler, 2000) database and reducing the redundancy of the original dataset with CD-HIT (Li and Godzik, 2006), we obtained 22,168 low-homologous enzyme sequences. These sequences are annotated with the Enzyme Commission (EC) system (Cornish-Bowden, 2014), which is a hierarchical classification system and the details can be referred to (Li *et al.*, 2018a). For the illustration purpose and without loss of generality, we used the first level labels, i.e., the six main classes of the EC system, for the experiments.

#### 4.1.2 2D HeLa Images

The HeLa image dataset^2^ (Boland and Murphy, 2001) contains the fluorescence microscope images of the 10 major subcellular structures in HeLa cells. Each image is a gray-scale image, whose dimensionality is 512 by 384. Based on the actual subcellular structure contained in the image, each image is labeled with one of the following 10 labels: ActinFilaments, Endosome, Endoplasmic Reticulum, Golgi gia, Golgi gpp, Lysosome, Microtubules, Mitochondria, Nucleolus, Nucleus. Within each class, there are roughly 90 images.

#### 4.1.3 BreakHis

The Breast Cancer Histopathological Database (Break His)^3^ (Spanhol *et al.*, 2016) is composed of 9,109 microscopic images of the breast tumor tissue, which were collected from 82 patients with four different magnifying factors (40X, 100X, 200X, 400X). Each image is a 3-channel RGB image, whose dimensionality is 700 by 460. These images are first classified into benign samples or malignant samples. The benign samples are further classified into four classes: adenosis (A), fibroadenoma (F), phyllodes tumor (PT), and tubular adenoma (TA). Similarly, malignant samples are also classified into four subclasses: carcinoma (DC), lobular carcinoma (LC), mucinous carcinoma (MC) and papillary carcinoma (PC). As a result, each image is annotated with one of the eight labels. For the experiments, we augmented this dataset to 20,696 by using the combination of images with different magnifying factors.

Notice that this dataset is unbalanced with the number of malignant images being two times as large as that of the benign images.

### 4.2 Compared Methods

We compared our method with five different methods. We refer the first method as the “All Data” method. When data from a new class appear, this method trains a deep learning model from scratch for multi-class classification, using all the new and old data. It can be expected that this method should have the highest classification performance. However, it is very computationally inefficient. In addition, the model trained with the new data can have completely different features from the one trained on the old data, which leads to poor model robustness and generalization (Parisi *et al.*, 2018). The second method is the iCaRL method (Rebuffi *et al.*, 2016), which is the state-of-the-art method for class incremental learning in the computer vision field. A brief introduction to iCaRL could be referred to Section 2.2. The third method is EWC, which is another recent work (Section 2.1). The fourth method is the “Fine Tune” method, in which we only use the new data to tune the model, without using any old data or the regularizers. The fifth method is the baseline “Random Guess” method, which assigns the label of each test data sample randomly without using any model.

### 4.3 Performance on EC Number Classification

As for this enzyme function prediction task, we first gave data of two EC classes to each method and trained them as a binary classifier. Then each time we incrementally gave data from one new EC class to each method, until all the six classes were fed to the model. Fig. 6(A)(B)(C) show the multi-class classification performance of the six methods, in terms of Kappa score, accuracy and Macro F1-score with respect to the number of classes.

**Fig. 6.**
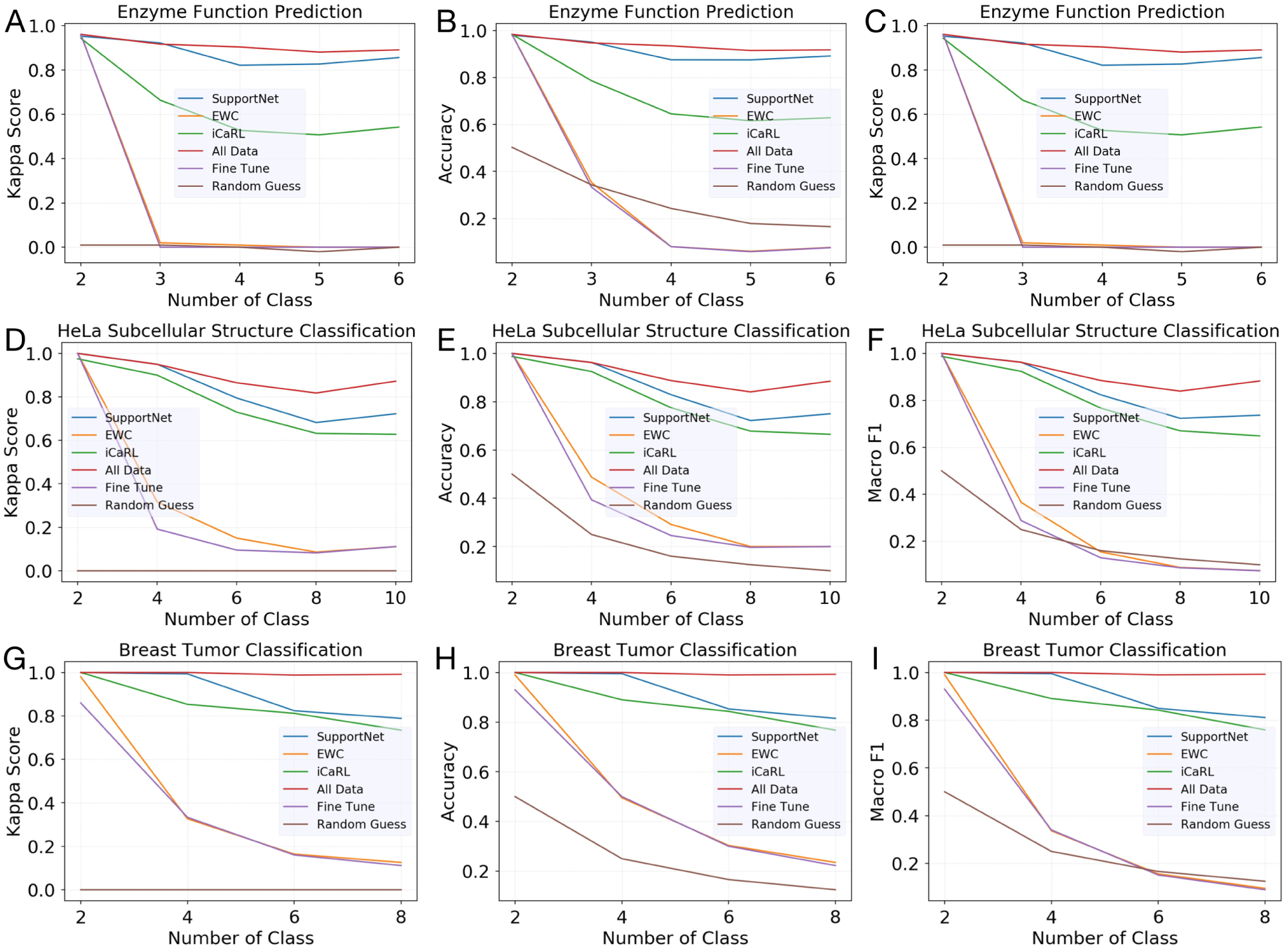
Performance comparison between SupportNet and five competing methods on the three tasks. For the SupportNet and iCaRL methods, we set the support data (examplar) size roughly one tenth of all the training data, that is, 2000 out of 20168 for the enzyme dataset, 80 out of 580 for the HeLa dataset, and 1600 out of 20296 (after augmentation) for the breast tumor dataset. (A)-(C): The Kappa score, accuracy, and macro F1-score of different methods over different numbers of classes on the enzyme function prediction task, respectively. (D)-(F): The Kappa score, accuracy, and macro F1-score of different methods over different numbers of classes on the subcellular structure classification task, respectively. (G)-(I): The Kappa score, accuracy, and macro F1-score of different methods over different numbers of classes on the breast tumor classification task, respectively. All the reported performance is over all the available classes at that time to the model.

As expected, the “All Data” method has the best classification performance because it trains a brand new model for each data set. However, it is an order of magnitude slower than our method and has poor model robustness because the model on the new data can be drastically different from the one on the old data. Nevertheless, the performance of the “All Data” method can be considered as the empirical upper bound of the performance of the incremental learning methods. Among the four incremental learning methods, they all have performance decrease to different degrees. EWC and “Fine Tune” have quite similar performance which drops quickly when the number of classes increases. Thus, although EWC has shown impressive performance on handling sequential tasks (Kirkpatrick *et al.*, 2017), it cannot handle bioinformatic tasks well, which is consistent with some previous reports (Parisi *et al.*, 2018; Kemker *et al.*, 2017). The iCaRL method is much more robust than these two methods. In contrast, SupportNet has significantly better performance than all the other incremental learning methods. In fact, its performance is quite close to the “All Data” method and stays stable when the number of classes increases. Specifically, the performance of SupportNet has less than 5% difference compared to that of the “All Data” method, yet is higher than the second best incremental learning method by at least 20%.

Another interesting finding is that although the “Fine Tune” method is much better than the “Random Guess” method on the binary classification task, its performance is even worse than that of the “Random Guess” method on multi-class classification. This is because that after the fine tuning, the model only focuses on the new data from the new class and forgets the knowledge learned from the old data, which illustrates the situation of catastrophic forgetting.

We further investigated the confusion matrix of the “Random Guess” method, the “Fine Tune” method, iCaRL and SupportNet (Fig. 7). As expected, the “Fine Tune” method only considers the new data from the new class, and thus is overfitted to the new class (Fig. 7(B)). The iCaRL method partially solves this issue by combining deep learning with nearest-mean-examplars, which is a variant of KNN (Fig 7(C)). SupportNet, on the other hand, combines the advantage of SVM and deep learning by using SVM to find the important support data, which efficiently preserve the knowledge of the old data, and utilizing deep learning as the final classifier. This novel combination can efficiently and effectively solve the incremental learning problem (Fig 7(D)).

**Fig. 7.**
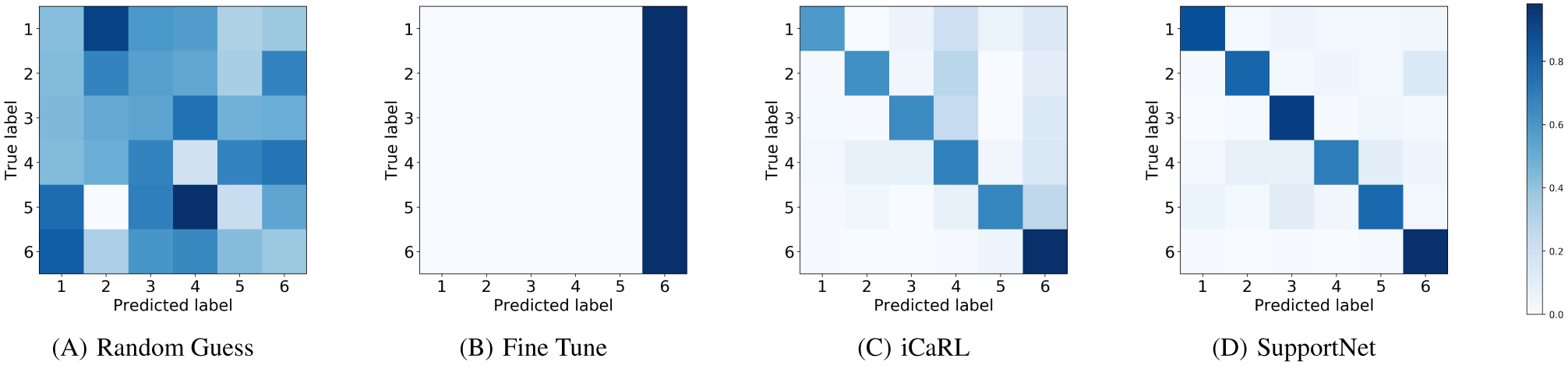
The confusion matrix of different methods on the 6-class classification task for EC prediction: (A) the “Random Guess” method, (B) the “Fine Tune” method, (C) iCaRL, and (D) SupportNet. The data from the first five classes were given as the old data, and the ones from the sixth class were given as the new data.

### 4.4 HeLa Subcellular Structure Classification and Breast Tumor Classification

For these two tasks, the basic experimental settings are similar to that of the first task, and the main difference is that we used data from two new classes as the new data during each round. The results of the HeLa subcellular structure classification task are shown in Fig. 6(D)(E)(F) and those of the breast tumor classification task are shown in Fig. 6(G)(H)(I). Similar conclusions could be drawn from these results: the “All Data” method performs the best and SupportNet is a close second. In addition, the results also demonstrate that although iCaRL was specifically designed for image classification in computer vision, SupportNet can still outperform it on these two image datasets, proving the effectiveness of our approach. Note that the two tasks are actually quite difficult for deep learning methods because of their small data size, with only 980 images belonging to 10 classes for the HeLa dataset and 9,109 original images belonging to 8 classes for the breast cancer dataset. Such small data size often results in overfitting for deep learning methods. In order to alleviate the overfitting problem in these tasks, we performed data augmentation with the augmentation ratio as 8, which is a common trick for deep learning methods to process images, before training these methods. As for the practical use of our method, we do not recommend training the model with data augmentation with a large augmentation ratio, because it may introduce too much noisy or even harmful information to the data. As an example, if we set the augmentation ratio as 2000 for the HeLa dataset, the performance for deep learning methods will significantly decrease.

### 4.5 Support Data Size

As reported by the previous study (Rebuffi *et al.*, 2016), the preserved dataset size can affect the performance of the final model significantly. We also investigated this problem in details here. As shown in Fig. 8, the performance degradation of SupportNet from the “All Data” method decreases gradually as the support data size increases, which is consistent with the previous study using the rehearsal method (Rebuffi *et al.*, 2016). What is interesting is that the performance degradation decreases very quickly at the beginning of the curve, so the performance loss is already very small with a small number of support data. That trend demonstrates the effectiveness of the support data selector in our framework, i.e., being able to select a small while representative support dataset. On the other hand, this decent property of our framework, which is guaranteed by the sparsity of support vector learning, is very useful when the users need to trade off the performance with the computational resources and running time.

**Fig. 8.**
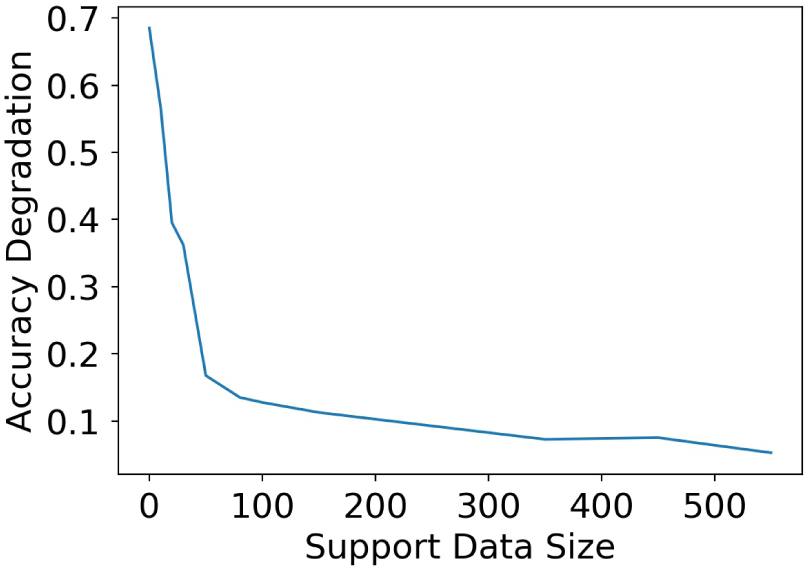
The accuracy degradation of SupportNet from the “All Data” method with respect to the size of the support data. The x-axis shows the support data size. The y-axis is the test accuracy degradation of SupportNet from the “All Data” method after incrementally learning all the classes of the HeLa subcellular structure dataset.

### 4.6 Regularizer Coefficient

Although the performance of the EWC method on incremental learning is not impressive (Fig. 6), the EWC regularizer plays an important role in our method. We investigated its influence in more details. We evaluated our method by varying the EWC regularizer coefficient from 1 to 100,000, and compared it with the “All Data” method and iCaRL (Table 1). We can find that the performance of SupportNet varies with different EWC regularier coefficients, with the highest one very close to the “All Data” method, which is the upper bound of all the incremental learning methods, whereas the lowest one having around 13% performance degradation. The results make sense because from the neurophysiological point of view, SupportNet is trying to reach the stability-plasticity balance point for this classification task. If the coefficient is too small, which means we do not impose enough constraint on those weights which contribute significantly to the old class classification, the deep learning model will be too plastic and the old knowledge tends to be lost. If the coefficient is too large, which means that we impose strong constraint on those weights even they are not important to the old class classification, the deep learning model will be too stable and does not have enough capacity to incorporate new knowledge. In general, our results are consistent with the stability-plasticity dilemma. Human beings, having been evolving for millions of years, have already found the right balance between the synaptic stability and plasticity, which enables human beings to deal with catastrophic forgetting well when learning new knowledge.

**Table 1.**
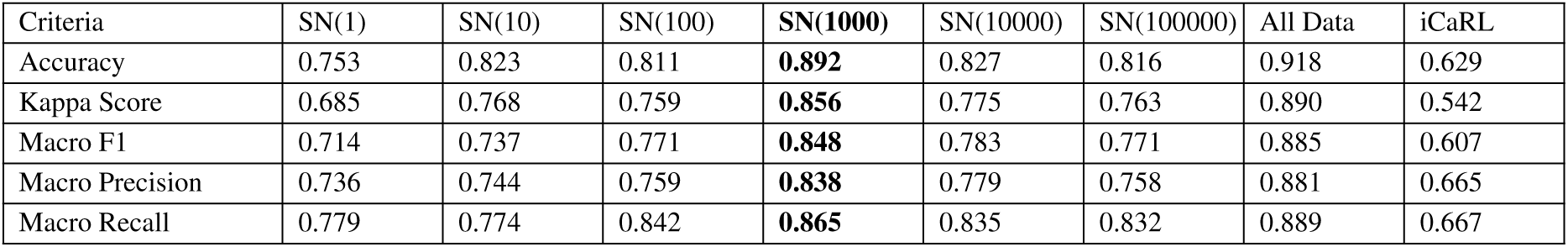
Performance of SupportNet with respect to different values of the EWC regularizer coefficient. The experiments were done on the enzyme function prediction task. All the results, except for the last two columns, are by incrementally learning all the six classes of the EC system one by one using different EWC regularizer coefficient values, with the support data size fixed to be 2,000. ‘SN’ stands for SupportNet. The numbers inside the bracket are the coefficient values. The last two columns show the performance of the “All Data” method and iCaRL with the examplar size as 2,000, respectively. The best performance of SupportNet is shown in bold.

| Criteria | SN(1) | SN(10) | SN(100) | SN(1000) | SN(10000) | SN(100000) | All Data | iCaRL |
| --- | --- | --- | --- | --- | --- | --- | --- | --- |
| Accuracy | 0.753 | 0.823 | 0.811 | **0.892** | 0.827 | 0.816 | 0.918 | 0.629 |
| Kappa Score | 0.685 | 0.768 | 0.759 | **0.856** | 0.775 | 0.763 | 0.890 | 0.542 |
| Macro F1 | 0.714 | 0.737 | 0.771 | **0.848** | 0.783 | 0.771 | 0.885 | 0.607 |
| Macro Precision | 0.736 | 0.744 | 0.759 | **0.838** | 0.779 | 0.758 | 0.881 | 0.665 |
| Macro Recall | 0.779 | 0.774 | 0.842 | **0.865** | 0.835 | 0.832 | 0.889 | 0.667 |

### 4.7 Underfitting and Overfitting

When training a deep learning model, one would encounter the notorious overfitting issue almost all the time. It is still the case for training an incremental learning model, but we find that there are some unique issues of such learning methods. Table 2 shows the performance of SupportNet and iCaRL on the real training data (i.e., the new data plus the support data for SupportNet and examplars for iCaRL), all the training data (i.e., the new data plus all the old data), and the test data. It can be seen that both methods perform almost perfectly on the real training data, which is as expected. However, the performances of iCaRL on the test data and all the training data are almost the same, both of which are much worse than that on the real training data. This indicates that iCaRL is overfitted to the real training data but underfitted to all the training data. As for SupportNet, the issue is much less severe than iCaRL as the performance degradation from the real training data to all the training data reduces from 37% as in iCaRL to 7% in SupportNet. This suggests that the support data selected by SupportNet can capture the distribution of all the training data accurately.

**Table 2.** Underfitting and overfitting of iCaRL and SupportNet. The experiments were done on the enzyme function prediction task. “Real training data” means the training accuracy on the new data plus the support data for SupportNet and examplars for iCaRL. “All training data” means the accuracy of the model trained on the real training data over the new data and all the old data. “Test data” means the accuracy of the model trained on the real training data over the test data.

| Method | SupportNet | iCaRL |
| --- | --- | --- |
| Real training data | 0.987 | 0.991 |
| All training data | 0.920 | 0.626 |
| Test data | 0.839 | 0.629 |

## 5 CONCLUSION

In this paper, we proposed a novel class incremental learning method, SupportNet, to solve the catastrophic forgetting problem by combining the strength of deep learning and support vector machines. SupportNet can identify the support data from the old data efficiently, which are fed to the deep learning model together with the new data for further training so that the model can review the essential information of the old data when learning the new information. With the help of two powerful consolidation regularizers, the support data can effectively help the deep learning model prevent the catastrophic forgetting issue, eliminate the necessity of retraining the model from scratch, and maintain stable extracted features between the old and the new data.

## ACKNOWLEDGEMENTS

This work was supported by the King Abdullah University of Science and Technology (KAUST) Office of Sponsored Research (OSR) under Awards No. FCC/1/1976-04, URF/1/2602-01, URF/1/3007- 01, URF/1/3412-01, URF/1/3450-01 and URF/1/3454-01. W.C. was supported by Basic Research Grant from Science and Technology Innovation Commission of Shenzhen Municipal Government [JCYJ20170 307105752508]. Y.H. was supported by the International Cooperation Research Grant (No. GJHZ20170310161947503) from Science and Technology Innovation Commission of Shenzhen Municipal Government.

## Footnotes

1 http://www.cbrc.kaust.edu.sa/DEEPre/dataset.html

2 http://murphylab.web.cmu.edu/data/2DHeLa

3 https://web.inf.ufpr.br/vri/breast-cancer-database/

